# Muscarinic acetylcholine receptors modulate HCN channel properties in vestibular ganglion neurons

**DOI:** 10.1101/2021.12.30.474193

**Authors:** Daniel Bronson, Radha Kalluri

**Affiliations:** Hearing and Communications Neuroscience Training Program, University of Southern California, Los Angeles, CA, 90057; Department of Otolaryngology-Head and Neck Surgery, Keck School of Medicine, University of Southern California, Los Angeles, CA, 90057; Zilkha Neurogenetic Institute, Keck School of Medicine, University of Southern California, Los Angeles, CA, 90057

## Abstract

Vestibular efferent neurons play an important role in shaping vestibular afferent excitability and, accordingly, on the information encoded by their spike patterns. Efferent- modulation is linked to muscarinic signaling cascades that affect ion channel conductances, most notably low-voltage gated potassium channels such as KCNQ. Here we tested and found that muscarinic signaling cascades also modulate hyperpolarization-activated cyclic-nucleotide gated channels (HCN). HCN channels play a key role in controlling spike-timing regularity and a non-chemical form of transmission between type I hair cells and vestibular afferents. The impact of cholinergic efferent input on HCN channels was assessed using voltage-clamp methods, which measure currents in the disassociated cell bodies of vestibular ganglion neurons (VGN). Membrane properties in VGN were characterized before and after administration of the muscarinic acetylcholine receptor (mAChR) agonist oxotremorine-M (Oxo-M). We found that Oxo-M shifted the voltage-activation range of HCN channels in the positive direction by 3.2 ± 0.7 mV, which more than doubled the available current when held near rest at -70 mV (a 139 ± 43% increase, n=12). This effect was not blocked by pre-treating the cells with a KCNQ channel blocker, linopirdine, which suggests that this effect is not dependent on KCNQ currents. We also found that HCN channel properties in the baseline condition and sensitivity to mAChR activation depended on cell size and firing patterns. Large-bodied neurons with onset firing patterns had the most depolarized activation range and least sensitivity to mAChR activation. Together, our results highlight the complex and dynamic regulation of HCN channels in VGN.

**Significance Statement:** Vestibular afferents express a diverse complement of ion channels. This diversity is key to shaping their various response properties, allowing them to encode both fast and slow head movements. *In vitro* studies that characterized afferent diversity identified low-voltage activated potassium channels and hyperpolarization-activated cyclic- nucleotide gated (HCN) channels as crucial for shaping the timing and sensitivity of afferent responses. Afferent excitability is known to be controlled by a network of acetylcholine-releasing efferent neurons that close a type of low-voltage activated potassium channel found on the afferent neuron. Here, we show that the same efferent signaling cascade that shuts these potassium channels also enhances the activation of HCN channels by depolarizing their voltage-activation range. The size of this effect varies from cell to cell depending on the endogenous properties of the HCN channel and on cell type (as determined by discharge patterns and cell size). Simultaneously controlling two ion-channel groups gives the vestibular efferent system robust and flexible control over the excitability and timing properties vestibular afferent activity.

## Introduction

The mammalian vestibular system detects head motions across a wide range of amplitudes and frequencies. Low and high-frequency head movements are encoded by two parallel streams of vestibular afferents that differ in excitability and spike-timing (Sadeghi et al., 2007; Eatock and Songer, 2011). Afferents that spike at regular intervals have lower detection thresholds for low-frequency head movements, while so-called irregular afferents have lower detection thresholds for all other frequencies. Afferent excitability and spike-timing reflect, in part, the complement of ion channel conductances they express. In vitro studies have identified low-voltage gated potassium channels (e.g., Kv1 and KCNQ) and hyperpolarization-activated cyclic-nucleotide gated (HCN) channels as critical regulators of afferent spike timing. These two ion channel groups have voltage- gated properties that make currents available between rest potential and voltage threshold, where they can influence the summation of synaptic events and action potential generation.

Vestibular efferents modulate afferent excitability through their impact on ion channels. Vestibular afferents receive input from an extensive network of primarily cholinergic efferent neurons, which increase afferent excitability by activating muscarinic acetylcholine receptors (mAChR). The resulting signaling cascades increase afferent excitability by shutting down low-voltage activated potassium currents carried by KCNQ channels (Pérez et al., 2009; Sadeghi et al., 2009; Holt et al., 2017; Raghu et al., 2019). Here, we consider whether muscarinic signaling cascades also affect HCN channels in vestibular afferent neurons. This is a possibility because the second messenger cascades initiated by mAChR activation include phosphatidylinositol 4,5-bisphosphate (PIP_2_) and cAMP (Zhang et al., 2003; Hughes et al., 2007); which are known modulators of HCN channels (Wainger et al., 2001; Pian et al., 2007). Therefore, we tested if HCN channels in vestibular afferents are sensitive to muscarinic signaling cascades.

Efferent modulation of HCN channels is significant because both channel density and voltage-activation properties impact vestibular afferent activity. HCN channels conduct an inward, hyperpolarizing-activated Na+/K+ current (*I*_H_) that famously controls pace-making in the heart and brain (Pape and McCormick, 1989; DiFrancesco, 1993; Biel et al., 2009). The *I*_H_ current has received significant attention in the vestibular system for its putative role in shaping spike-timing regularity (Horwitz et al., 2014; Yoshimoto et al., 2015). HCN channels are also thought to mediate a non-chemical form of communication (Yamashita and Ohmori, 1991; Songer and Eatock, 2013; Highstein et al., 2014) by resistively coupling sensory hair-cells to vestibular afferent neurons (Contini et al., 2020). However, HCN channels can only shape neuronal responses when the HCN current, I_H_, is available between rest and threshold. The HCN channel is highly modifiable, so characterizing how HCN channels are regulated in the vestibular system remains an important goal.

This study extends our understanding of HCN channel properties in vestibular afferent neurons under experimental conditions that preserve endogenous signaling cascades and intracellular second messengers. To do this, we recorded from vestibular ganglion neurons (VGN; the cell bodies of vestibular afferents) using the perforated-patch clamping technique to avoid disturbing the intracellular signaling cascades typically disrupted by standard whole-cell patch-clamping methods. This further allowed us to test if HCN channels are sensitive to the intracellular signaling triggered by mAChR activation in these neurons. The recordings were made in VGN older than post-natal day (P)9 to allow for the developmental upregulation of both HCN and KCNQ channels; the latter are the primary ion channel target for muscarinic signaling cascades.

Our findings reveal a previously unrecognized diversity in the activation properties of HCN channels that is related to both cell size and firing pattern. Large, transient-firing VGN have HCN channels that activate at more positive potentials than do the medium to small-sized VGN. Moreover, mAChR signaling cascades depolarize HCN activation range in VGN neurons. There was also a size dependence; HCN channels were most sensitive to mAChR activation in small neurons. Our results paint a rich and complex picture of how HCN channel properties vary and dynamically change in vestibular ganglion neurons.

## Materials and Methods

### Preparations

Data were recorded from the superior portion of the vestibular ganglion of Long- Evans rats of either sex aged post-natal day (P)9-18 (P0, birth day). All animals were handled and housed in accordance with National Institutes of Health *Guide for the Care and Use of Laboratory Animals.* All animal procedures were approved by the University of Southern California Institutional Animal Care and Use Committee. Chemicals were obtained from Sigma-Aldrich unless otherwise specified. The temporal bones from the animals were dissected in chilled and oxygenated Leibowitz medium supplemented with 10 mM HEPES (L-15 solution). The superior part of the vestibular ganglia was detached from the distal and central nerve branches and separated from the otic capsule. Bone fragments, debris and any remaining connective tissue were removed from the surface of the ganglia. Ganglia from 2 litter-matched animals of either sex were pooled together. Ganglia were then incubated at 37°C in L-15 medium solution with 0.05% collagenase and 0.25% trypsin for 20-40 min, depending on age of the animal. The ganglia were then washed in fresh L-15 solution, dissociated by triturating through a series of polished Pasteur pipettes and allowed to settle onto 1% polyethyleneimine-coated glass coverslips. Culture dishes contained bicarbonate- buffered culture medium (minimal essential medium, Invitrogen), supplemented with 10 mM HEPES, 5% FBS, and 1% penicillin-streptomycin (Invitrogen) . The culture medium was titrated with NaOH to a pH of 7.35. Cells were incubated for 16-24 hrs in 5% CO_2_/95% air at 37°C. Short-term incubation tends to remove supporting and satellite cells, clears debris from enzyme treatment, allows cells to adhere to the substrate, and promotes successful recordings as the animals entered the second post-natal week and beyond.

### Electrophysiology

Cells were viewed at 40x using an inverted microscope (Zeiss, Axiovert 135 TV) fitted with Nomarski optics. A MultiClamp 700B amplifier, Digidata 1440 board, and pClamp 10.7 software (MDS; RRID: SCR_011323) were used to deliver, record, and amplify all signals. Recording pipettes were fabricated using filamented borosilicate glass. Pipettes were fire polished to yield an access resistance between 4 and 8 MΩ. Recording pipettes were coated with parafilm (Bemis Company Inc) to reduce pipette capacitance.

The properties of ion channels were studied using perforated-patch methods. The contents of the perforated-patch internal solution contained the following (in mM): 75 K_2_SO_4_, 25 KCl, 5MgCl_2_, 5 HEPES, 5 EGTA, 0.1 CaCl_2_, and titrated with 1 M KOH to a pH of 7.4 and an osmolality of ∼260 mmol/kg. Amphotericin B (240/ml; Sigma-Aldrich) was dissolved in DMSO and added to the perforated-patch solution on day of recording. This allowed passage of small monovalent ions while preventing larger molecules from dialyzing.

The series resistance was estimated using whole-cell compensation in voltage-clamp and ranged between 8 and 35 MΩ. Although series resistance tends to be higher in perforated patch than ruptured patch, the same cutoff value of 35 MΩ has been used in our previous studies (Ventura and Kalluri, 2019). Series resistance was left uncompensated and instead corrected off-line. In a subset of experiments, we applied 50% series resistance compensation to better understand the influence of series resistance to our *I*_H_ measurements and found that off-line compensation produced similar values and that series resistance was not a significant predictor of *I*_H_ activation (data not shown).

Recordings were made at room temperature (25-27°C) and in an external bath continuously perfused with fresh oxygenated L-15 media. Perforated patch internal solution had junction potentials of +5.0 mV, which was computed with JPCalc (Barry, 1994) as implemented by pClamp 10.7 and left uncorrected. Only recordings in which the cell had formed a gigaOhm seal were used. Recording stability was monitored by measuring resting potential, input resistance, series resistance and size of action potential and significant fluctuations in these values within each experimental condition was indicative of an unhealthy cell and/or compromised recording and was not included in our dataset.

### Pharmacology

10 or 100 µM solutions of the mAChR agonist oxotremorine-M (Oxo-M) were prepared on the day of the experiment by dilution in L-15. The effect of Oxo-M on *I*_H_ was equally saturated at both doses (Average shift in *I*_H_ *V*_1/2_ with 10 µM Oxo-M = 4.4 ± 0.8, n=4, 100 µM = 4.0 ±1.2, n=17, student t-test t(20)=0.16, p=0.8754, n=21).

Therefore, we pooled both concentrations for our statistical analyses. Figures include Oxo-M concentration whenever relevant. In the experiments that used linopirdine, a 10 µM solution was prepared on the day of the experiment by dilution in L-15. Drugs were applied via pressurized super-perfusion system (Warner Instruments). Measurements of *I*_H_ were made at least 5 min after exposure to any drug.

### I_H_ activation parameters

We studied the voltage-dependent activation of *I*_H_ using a voltage-clamp tail- current protocol (Fig. 2). The protocol begins with a pre-conditioning voltage step, then a 100 mV “tail-step” for 300 ms and finally returns to -60 mV for 100 ms. The pre- conditioning voltage step is incremented by 5 mV to range between -135 and -40 mV. A 15 s delay was implemented between each step to allow enough time for HCN channels to close completely before starting the next voltage step.

The HCN channel that carries I_H_ opens very slowly and the voltage-dependent activation of *I*_H_ is sensitive to the length of the pre-conditioning hyperpolarization. We chose to make all measurements with a 1.7s pre-conditioning step. Although even longer steps would allow more current to activate and reach its true steady state, the extreme hyperpolarization significantly increases the cell’s risk of dying during recording (Ventura and Kalluri, 2019). Therefore, we chose the shorter duration step to allow us to repeat the protocol in several pharmacological conditions.

The voltage-dependent activation of *z*_H_ is characterized by plotting the currents flowing during the tail step (*I*_tail_) against the pre-condition voltage (*V*_pre_). These data were fit with a Boltzmann function (Eq. 1):

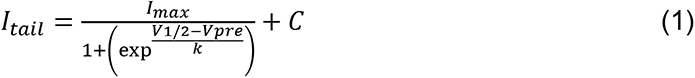

where *I*_max_ is the maximum current measured during the tail, *V*_pre_ is the pre-conditioning / test voltage, *V*_1/2_ is the half-activation potential, *k* is the slope factor, and *C* is a constant. Measurements of *I*_tail_ were taken 100 ms into the tail step to avoid contamination from other low-voltage-gated currents, particularly during the most positive voltage steps (-60 to -40mV).

The time course for *I*_H_ current activation was estimated by fitting the current evoked at the -135 to -70 mV pre-conditioning step with a single exponential. All fits were obtained after a 60 ms delay from the start of the voltage step. Only fits with a correlation coefficient ≥ 0.95 were retained. The *τ* value was used to estimate the time- course of *I*_H_ activation. Values greater than 5 ms, which were often observed at the smallest voltage steps (± 10 mV), were discarded since these values greatly exceeded our 1.7 s *I*_H_-testing protocol and cannot be assumed to be accurate. To estimate the effect of Oxo-M on the voltage activation speed of HCN channels, the *τ* values across multiple voltage steps were fitted with a simple exponential equation shown in Equation 2. This fit allowed us to account for the positive shift in voltage-activation range caused by Oxo-M and quantify the extent to which the current activated fully at smaller voltage steps. Only fits with a correlation coefficient ≥ 0.80 are included in figures and our analyses. From this equation, we quantified the slope of the relationship between *τ* and voltage-step as a single model parameter, α.

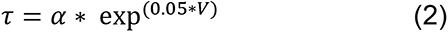

### Analysis

*General*. All data were analyzed with pClamp 10 software (Clampfit; MDS Analytical Technologies). Input resistance (*R*_in_) was calculated from voltage changes in response to a hyperpolarizing 10 or 20 pA step in current-clamp mode. The membrane time constant (*τ*_m_) was assessed by fitting a single exponential to the current transient measured in response to a 5 mV depolarizing voltage step in voltage-clamp mode.

Membrane capacitance (*C*_m_) was calculated by dividing *τ*_m_ by the series resistance (*R*_s_). On-line calculations of *C*_m_ were verified off-line using the average of multiple depolarizing steps recorded with whole-cell compensation turned off.

Statistical analysis was done with Origin Pro (OriginLab; RRID: SCR_014212) and/or JMP Pro 13 (SAS Institute; RRID:SCR_014242). Statistical significance was estimated with Student’s t test when variances were equal or with Welch’s t test when variances were not equal. We used an ɑ level of 0.05 for all statistical tests. One-way and two-way ANOVAs were applied followed by a post hoc Tukey’s HSD analysis, as required, to drug condition and firing pattern on I_H_ half-activation. Statistics are reported as means ± SEM.

## Results

We present whole-cell, perforated-patch clamp recordings from cultured VGN of rats aged post-natal day (P)9 to P22 (n = 81). Firing pattern was assessed in all recorded cells, and *I*_H_ was characterized in 48 of the 81 cells recorded in this study.

First, we describe the diversity of HCN current characteristics in VGN and its sensitivity to recording conditions. Second, we test whether HCN channels in VGN are sensitive to muscarinic signaling cascades using the muscarinic agonist Oxotremorine-M (Oxo-M). Third, we characterize HCN channel sensitivity to muscarinic signaling cascades in relation to cell type as defined by firing pattern and somatic size. Finally, we examine the impact of muscarinic signaling cascades on firing patterns. Table I summarizes the number of cells included in each analysis.

**Table 1.**

| **Data Collected** | **Control (n)** | **Linopirdine (n)** | **Oxo-M (n)** | **Linopirdine + Oxo-M (n)** |
| --- | --- | --- | --- | --- |
| **Firing Pattern** | Total n = 81<br>Transient (n=40),<br>Sustained-A (n=5),<br>Sustained-B (n=26),<br>Sustained-C (n=10) | Total n = 10<br>Transient (n=5)<br>Sustained-A (n=1),<br>Sustained-B (n=3),<br>Sustained-C (n=1), | Total n = 21<br>Transient (n=13)<br>Sustained-B (n=1),<br>Sustained-B (n=6),<br>Sustained-C (n=1), | Total n = 12<br>Transient (n=4)<br>Sustained-A (n=1),<br>Sustained-B (n=4),<br>Sustained-C (n=3), |
| **$I_H$ Voltage Activation Range** | Total n = 48<br>Transient (n=29),<br>Sustained-A (n=1),<br>Sustained-B (n=11),<br>Sustained-C (n=7) | Total n = 10<br>Transient (n=5)<br>Sustained-A (n=1),<br>Sustained-B (n=3),<br>Sustained-C (n=1), | Total n = 21<br>Transient (n=13)<br>Sustained-B (n=1),<br>Sustained-B (n=6),<br>Sustained-C (n=1), | Total n = 12<br>Transient (n=4)<br>Sustained-A (n=1),<br>Sustained-B (n=4),<br>Sustained-C (n=3), |
| **HCN Channel Activation Rate** | Total n = 22<br>Transient (n=14)<br>Sustained-B (n=5),<br>Sustained-C (n=3), | Total n = 3<br>Transient (n=1)<br>Sustained-B (n=2)<br>*data not shown* | Total n = 16<br>Transient (n=10)<br>Sustained-A (n=1)<br>Sustained-B (n=5) | Total n = 6<br>Transient (n=2)<br>Sustained-A (n=1)<br>Sustained-B (n=3) |

### Diversity in the activation properties of HCN channels in the vestibular ganglion

We quantified the voltage-sensitive properties of *I*_H_ using a voltage-clamp tail protocol consisting of 1.7-s pre-conditioning steps from -135 to -35 mV in 5 mV increments followed by a tail step to -100 mV (Figure 1A). The goal of the 1.7-s pre- conditioning step was to activate the slowly-activating hyperpolarization-gated inward currents conducted through the HCN channels. We assessed two properties of HCN channel activation that are known to be modifiable: voltage sensitivity and rate of activation.

**Figure 1.**
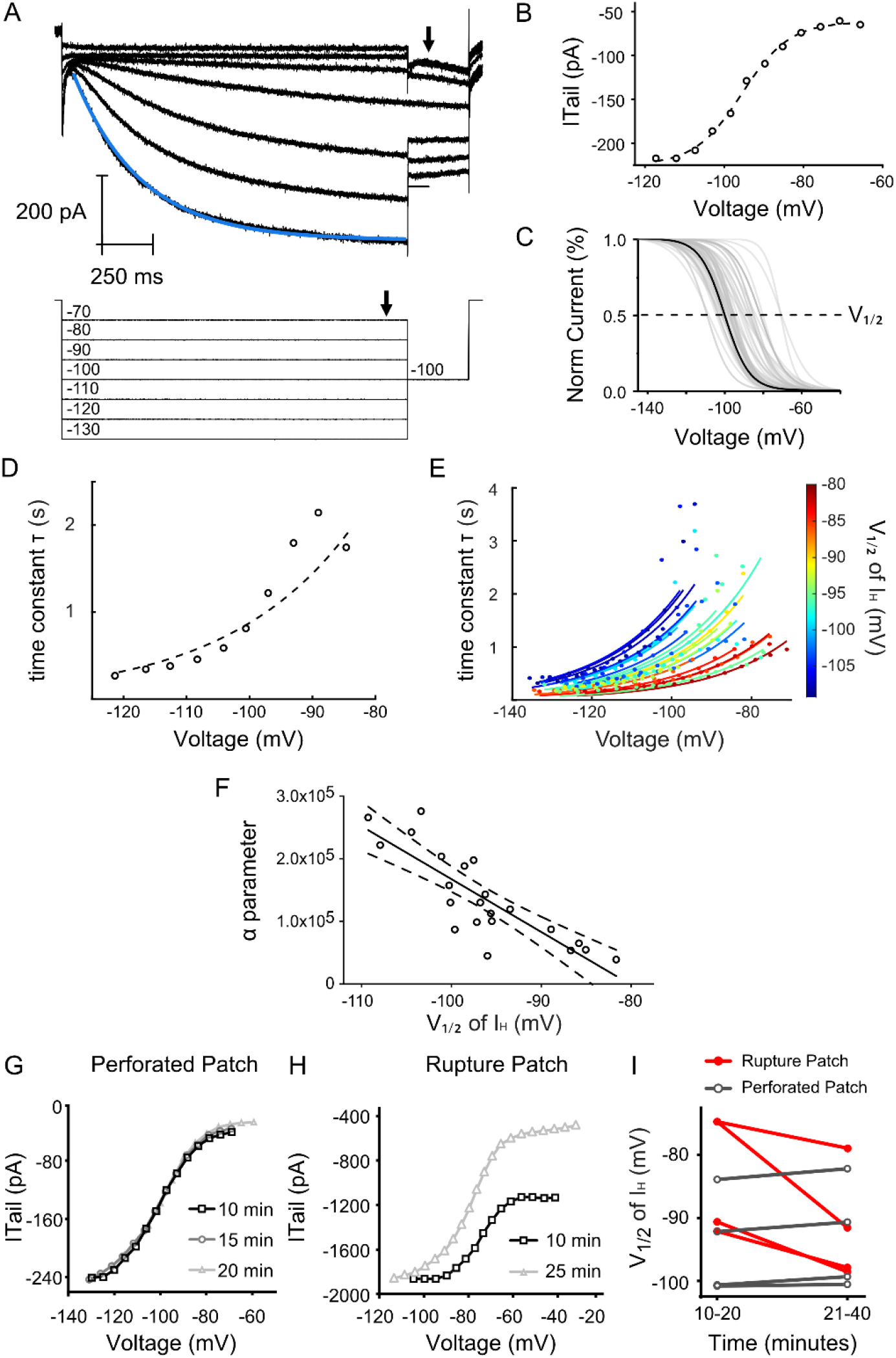
VGN have HCN channels with diverse voltage-gated activation properties. **A.** Currents activated from voltage-steps from -130 to -70 mV in a single cell. A single-exponential fit in blue is overlaid on the current response to each pre- conditioning step (example exponential fit is shown in blue) **B.** *I*_H_ activation measured as the magnitude of the tail current (arrow head in A, top) as a function of the pre- conditioning voltage step (arrow head in A, bottom). *I*_H_ activation curves were fit by Boltzmann function from Eq. 1 of the methods (dotted curve) **C.** *I*_H_ activation curves fit by Boltzmann function are then normalized to the maximum conductance of *I*_H_. Gray curves show the activation curves measured in 48 cells, bold curve shows the activation curve for cell shown in A and B. **D.** Current responses were fit with an exponential line (Exponential fit of current response to -130 mV step is shown in blue in A). The time constant *τ* from the exponential fits of the cell shown in A are plotted with the voltage- step on the x-axis. A single-parameter exponential described in Eq. 2 of the methods was used to fit the τ value / voltage relationship in this cell (dotted line). **F.** Fits of the τ value / voltage relationship for each cell. Points and lines are color-coded according to the *V*_1/2_ of each individual cell, with the most depolarized *V*_1/2_ values in red and the most hyperpolarized *V*_1/2_ values in blue. **J.** The *α*parameter, which describes the steepness of the fits in *I*_H_ activation rate, are shown for each cell plotted against the *V*_1/2_ of *I*_H_.**G.** Tail currents from a single neuron assessed at three different time points using the perforated patch technique. **H.** Tail currents from a different single neuron assessed at two different time points using the rupture patch technique. **I.** Shift in the half-activation voltage of I_H_ (*V*_1/2_) after short (10-20 mins) and long (21-40) intervals.

The voltage dependence of *I*_H_ activation is apparent when the magnitude of the current measured during the tail step (-100 mV, arrow in Figure 1A top) is plotted as a function of the pre-conditioning voltage step (arrow in Figure 1A bottom, circles in Figure 1B). These current-voltage curves are sigmoidal in shape and fit a Boltzmann equation (dashed line in Figure 1B, see Equation 1 in methods). *I*_H_ activation curves derived from this protocol closely approximate those extracted by pharmacologically isolating *I*_H_ using HCN channel blockers like CsCl or ZD7288 (Ventura and Kalluri, 2019). To ensure adequate fitting, the stimulus range was adjusted in each cell to ensure that the *I*_H_ current saturated at the most negative potentials. Only Boltzmann curves with correlation coefficients of 0.95 or greater were retained to ensure that the voltage-activation properties were well described by the fit. The size of the maximum *I*_H_ current ranged from as large as 807 pA to as small as 33 pA (mean = 258.9 ± 24.0 nA, SD = 166.4 n= 48). The maximum size of the *I*_H_ current (*I*_max_) varied with membrane capacitance, which is an indirect measure of cell size (Limón et al., 2005). We found that capacitance is bimodally distributed and we classified eleven VGN as large according to previously established criteria of greater than 30 pF in capacitance (*C*_m_) (Almanza et al., 2012). Large VGN had larger maximum *I*_H_ currents than small VGN (large VGN maximum I_H_ current = 449.9 ± 62.7 pA, n=11 small VGN maximum *I*_H_ current = 202.0 ± 16.1 pA, n=37, t(47)=14.6, p<.0001, total n=48). However, maximum current density (*I*_max_/*C*_m_) was the same in both groups (t(47)=0.16, p=0.88, n=48) with an average of 12.0 ± 0.81 pA/pF in all cells recorded. To account for the variability in the size of *I*_H_ currents, we normalized the current-voltage curves using Boltzmann fits to compare the activation range from different cells (Figure 1C, example from Figure 1A bolded). The voltage at which half of the *I*_H_ current is active (V_1/2_) is indicated in Figure 1C as a dotted line. V_1/2_ ranged from as positive as -70 mV to as negative as -109 mV (Average V_1/2_ = -95.5 ± 1.1 mV, n=48).

The *I*_H_ activation rate depends on the size of the pre-conditioning voltage step with more negative steps activating the current more quickly than more positive steps. To quantify the relationship between voltage and activation rate, we fit a single exponential curve with a time constant (*τ*) to the *I*_H_ trace during each of the long pre- conditioning voltage steps (*V*) (blue curve overlaid on current response to -130 mV hyperpolarizing step in Figure 1A). *τ* measurements were discarded if they greatly exceeded the 1.7 s pre-pulse duration as they could not be assumed to be accurate (see *IH activation parameters* in *Methods*). Figure 1D shows the relationship between *τ* and the command voltage-steps, *τ*(*V*), in the same cell shown in Figure 1A. *τ* values were smallest (i.e., *I*_H_ activated more quickly) in response to the largest hyperpolarizing steps while *τ* values were largest at the smallest pre-conditioning steps (± 10 mV). We quantified the sensitivity of the *I*_H_ activation rate to voltage by fitting *τ*(*V*) with an exponential function with one free parameter, α (see Eq. 2 in Methods). α reflects the steepness of *τ*(*V*) and therefore how quickly the channel activates across a range of membrane potentials. In Figure 1D, the fit for the example cell is shown in the dashed line. The exponential fits of *τ*(*V*) show that the rate of activation is voltage sensitive is fastest (and *τ* is smallest) in response to the most hyperpolarized steps and is slower (*τ* is larger) in response to pre-conditioning steps near or more positive than the half-activation voltage. At potentials positive of *V*_1/2_, the reduced size of the *I*_H_ current and emergence of other currents makes it difficult to accurately characterize the kinetics of *I*_H_ in response to these voltage steps. Therefore, steps positive of -70 mV are excluded from activation rate analyses (see *I_H_ activation parameters,* Materials and Methods).

We assessed *I*_H_ voltage-sensitivity and whether VGN with more depolarized activation ranges also have faster activation kinetics. Other studies have linked HCN channel activation rate and voltage activation range since increased cAMP both shifts the voltage-activation range in the depolarizing direction and increases the rate of activation in VGN (Almanza et al., 2012; Ventura and Kalluri, 2019). *τ*(*V*) varied from cell to cell; both in the maximal activation rate of *I*_H_ and in its voltage-dependence (Figure 1E, n=22). Figure 1E color-codes exponential fits according to the half-activation voltage of *I*_H_ (V_1/2_). and cells with more depolarized *V*_1/2_ (red) have smaller values for τ(*V*) and faster activation rates than cells with hyperpolarized *V*_1/2_ (blue). To quantify the covariance between *V*_1/2_ and activation kinetics, we plotted *V*_1/2_ against the single parameter of our exponential fit, α (Figure 1F, see Eq. 2 in Methods). Larger values of α correspond to *τ* values that more quickly increase (i.e., slower activation) as the pre-conditioning steps approach the half-activation voltage. As illustrated in Figure 1F, α is inversely dependent on *V*_1/2_, which shows that HCN channel activation kinetics are faster in cells that have a depolarized *I*_H_ activation range.

There was significant cell-to-cell variability in the half-activation voltage (*V*_1/2_), which was broadly distributed between -70.4 mV and -109.3 mV (mean = -95.5 ± 1.1 mV, SD = 7.4 mV, n=48). The range of *V*_1/2_ values seen here is broader than in previous studies (Almanza et al., 2012; Ventura and Kalluri, 2019). These studies mostly recorded cells using rupture-patch methods in which the intracellular composition is subject to dialysis with the contents of the recording pipette. This dialysis may reduce endogenous cell-to-cell variability since HCN channels are sensitive to the intracellular of second messengers such as cAMP. Figure 1G and 1H illustrate the stability of *V*_1/2_ in an individual cell during recordings using perforated patch as compared to the instability of *V*_1/2_ in a second cell using ruptured patch methods. In perforated patch, the estimates of *I*_H_ currents are stable over the duration of the recording session, both in current magnitude and in *V*_1/2_ (Figure 1G). In contrast, there is considerable drift over time in both the current magnitude and *V*_1/2_ when recordings were made in rupture-patch (Figure 1H). In less than an hour, all four cells recorded using the ruptured patch technique experienced a hyperpolarizing shift in the *I*_H_ activation range, while the *I*_H_ activation range remained stable in all cells recorded using the perforated-patch technique (Figure 1I). The remaining data are reported only for recordings made using perforated-patch methods.

### Activating muscarinic acetylcholine receptors depolarizes the activation range of HCN channels via a KCNQ-independent mechanism

To test if activating muscarinic receptors influences the voltage-gated properties of HCN channels, we measured the voltage activation of *I*_H_ currents under different pharmacological conditions (Figure 2A and Figure 2B). First, we characterized *I*_H_ in a control solution (Figure 2A.1 and Figure 2B.1). In some cells, we switched the bath solution to contain Oxotremorine (Oxo-M; Fig. 2A.2), a muscarinic receptor agonist known to close KCNQ channels in VGN by activating a G-protein coupled signaling cascade. In other cells, we first added a KCNQ channel blocker, linopirdine (Lino; Fig. 2B.2) before switching to a cocktail containing both Lino and Oxo-M (Lino+Oxo-M; Fig. 2B.3). In order to maximize the number of successful recordings, we continued recording from new cells after Lino, Oxo-M, or Lino+Oxo-M were introduced into the bath. In these not-naïve cases, a control-condition recording was not available for a paired comparison before and after treatment.

**Figure 2.**
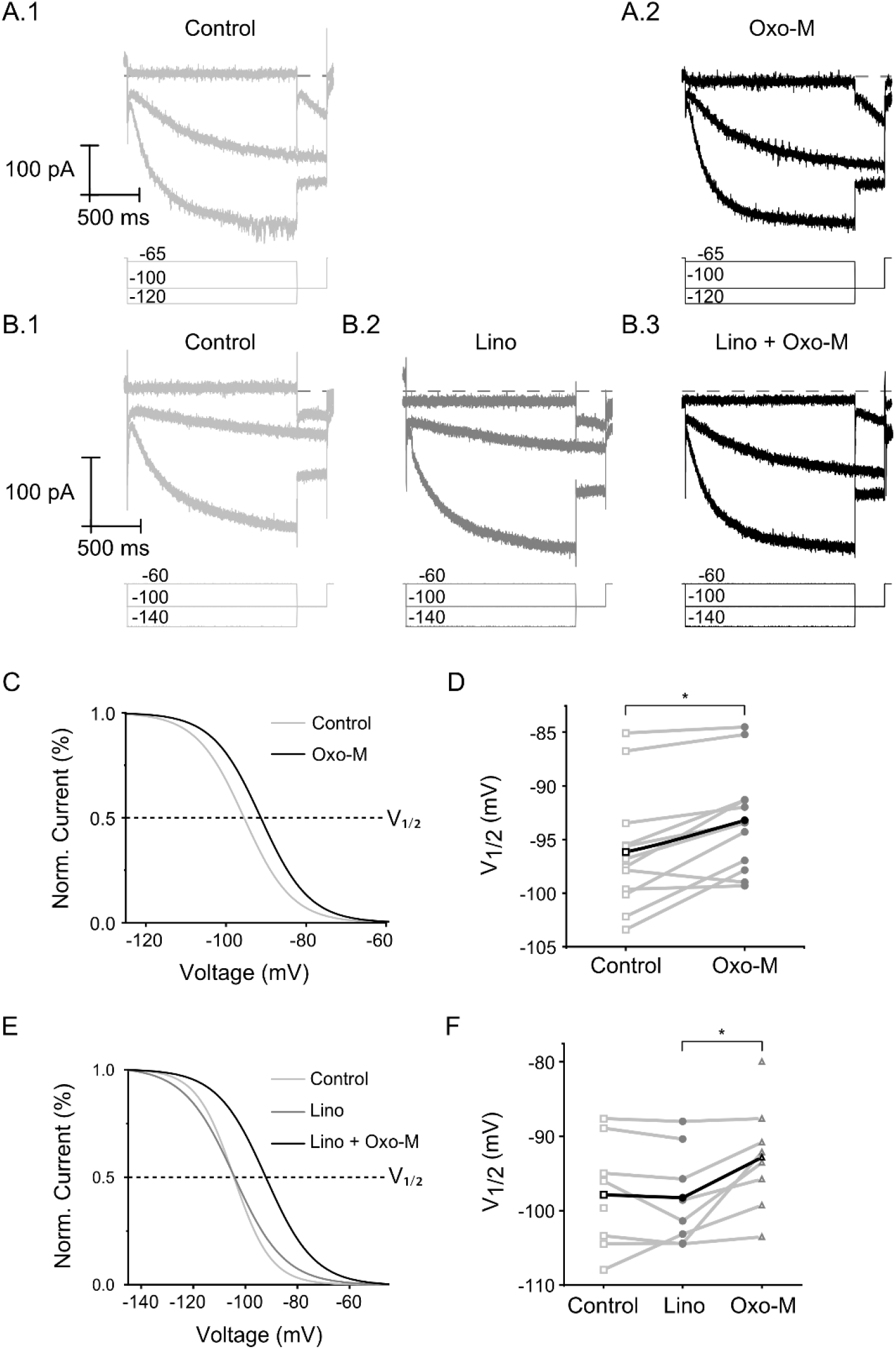
Activation of muscarinic acetylcholine receptors depolarizes the voltage-activation range of *I*_H_. **A.** Currents activated from voltage-steps from -135 to - 60 mV in a single before and after Oxo-M (A.1 and A.2) and in a single cell in control, linopirdine, and linopirdine and Oxo-M conditions (B.1, B.2, and B.3). C and E show data from the individual neuron shown in A and B, respectively. **C.** *I*_H_ activation curves were fit by Boltzmann function and the fractional activation of *I*_H_ plotted as a function of the pre-conditioning voltage step before (light gray) and after Oxo-M (black). **D.** Each point represents a single *V*_1/2_ measurement with lines connecting each cell before (square) and after Oxo-M (circle). The average shift of all cells is drawn in bold. V_1/2_ in 10 of 12 neurons shifted in the depolarized direction (* = p = 0.0018, paired t-test). **E.** I_H_ activation curves were fit by Boltzmann function and the fractional activation of *I*_H_ plotted as a function of the pre-conditioning voltage step in control (light gray), linopirdine (dark gray) and after Oxo-M (black). **F.** Each point represents a single *V*_1/2_ measurements at baseline (square), after linopirdine (circle) and then after administration of both linopirdine and Oxo-M (triangle). The average shift of all cells is drawn in bold. Linopirdine had no effect on V_1/2_, while the cocktail containing linopirdine and Oxo-M shifted the activation range in the depolarizing direction (* = p = .0324, Tukey’s HSD).

Activating the muscarinic receptor via Oxo-M enhanced HCN channels by shifting the activation range of *I*_H_ in the depolarizing direction. Figure 2C shows the voltage-activation of the *I*_H_ current in control and drug conditions for the cell shown in 2A. The *I*_H_ half-activation voltage, V_1/2_, is shown as a dotted line in Figure 2A. The voltage-dependent activation of *I*_H_ shifted in the depolarizing direction in 10 of the 12 cells tested, with one cell showing no change and another exhibiting a slight hyperpolarizing shift (Figure 2D). The extent of the depolarizing shift was variable from cell to cell; some cells saw a dramatic shift of as much as 6.3 mV while others had more modest shifts. The overall effect on *I*_H_ was a significant positive shift of 3.2 ± 0.7 mV (Paired t-test, t(11)=4.78, p = 0.0006, n=12). This depolarizing shift was consistent across the whole voltage range as the steepness of the fractional activation of *I*_H_ (the slope factor, k) was unaffected (k_control_ = 6.56 ± 0.37, k_Oxo-M =_ 6.44 ±0.18 mV, t(11)=0.32, p=0.753, paired t-test). Therefore, Oxo-M induced a depolarizing shift in the voltage activation range while the shape of the *I*_H_ current’s voltage sensitivity remained consistent.

We explored the mechanism of muscarinic enhancement of the *I*_H_ current by characterizing the involvement of the primary target of muscarinic signaling cascades in VGN: KCNQ channels. Activation of mAChR is known to reduce potassium currents carried by KNCQ channels (Pérez et al., 2009; Brown, 2010; Holt et al., 2017). To test if directly blocking the KCNQ channels had a similar impact on the activation of HCN channels as did the muscarinic agonist, we characterized *I*_H_ activation with the KCNQ channel blocker linopirdine in the bath. Figure 2E shows the voltage-activation of the *I*_H_ current in control and drug conditions for the cell shown in 2B. We verified that Linopirdine closed KCNQ channels by recording the change in the resting membrane potential, which depolarized by 5.7 ± 1.2 mV (t(8)=19.6, p = 0.0022, paired t-test, n=9). The activation range of *I*_H_ did not change significantly after blocking KCNQ with Linopirdine (Δ*V*_1/2_ = 0.61 ± 1.1 mV in the hyperpolarizing direction; t(6)=0.61, p = 0.61, paired t-test, n=7; Figure 2D). While Linopirdine-alone had no effect on *V*_1/2_, the addition of Oxo-M shifted *I*_H_ in the depolarizing direction (F(2,13)=5.86, p=0.0149, repeated measures ANOVA, post-hoc Tukey HSD, p = .0324) (Figure 2F). These results show that Oxo-M shifts *I*_H_ activation in the positive direction independently of its effect on closing KCNQ channels. These results are consistent with the idea that mAChR activation initiates second messenger cascades (e.g., cAMP or PIP_2_ ) that depolarize the activation range of HCN channels. It remains to be determined which second messengers are responsible for this effect. Since we found that linopirdine has no effect on HCN channels, cells that received both linopirdine and Oxo-M are included in the Oxo-M group unless otherwise noted.

### mAChR agonists increase the speed of HCN channel activation

We next examined whether muscarinic signaling cascades affect the activation rate of HCN channels. Since cells with more depolarized *V*_1/2_ also have faster activation kinetics (Figure 1F), we assumed that muscarinic signaling cascades would increase the rate of activation. However, variable expression of HCN channel isoforms that control activation kinetics and cAMP sensitivity suggests that mAChR-responsivity vary among vestibular afferents (Almanza et al., 2012). Therefore, variability in the sensitivity of the activation rate to muscarinic signaling cascades may reflect endogenous heterogeneity in HCN channel isoform and / or second messenger concentration within vestibular afferents.

Figure 3 shows evoked currents in a single cell before (Fig. 3A.1) and after Oxo-M (Fig. 3A.2). To characterize the activation time course in each condition, we fit single exponential curves to the currents evoked by voltage steps from -140 to -70 mV in 32 cells before and after adding Oxo-M to the bath. The time constants of these exponential fits are shown as a function of command voltage in Figure 3C. τ(V) and exponential fits to τ(V) in control condition are drawn with blue symbols and lines, respectively. Red symbols and curves are for the Oxo-M condition. τ(V) is generally smaller across all voltage steps (V) and is less sensitive to changes in voltage in Oxo-M (red curves) than in control condition (blue curves) (Figure 3C). This trend is quantified by comparing the parameter *α*from the fits to *τ*(V). Oxo-M reduced *α* values by 21.4 ± 4.9%, which reflects a significant increase in activation rate (t(10)=-2.49, p=0.0322, N=11, paired t-test).

**Figure 3.**
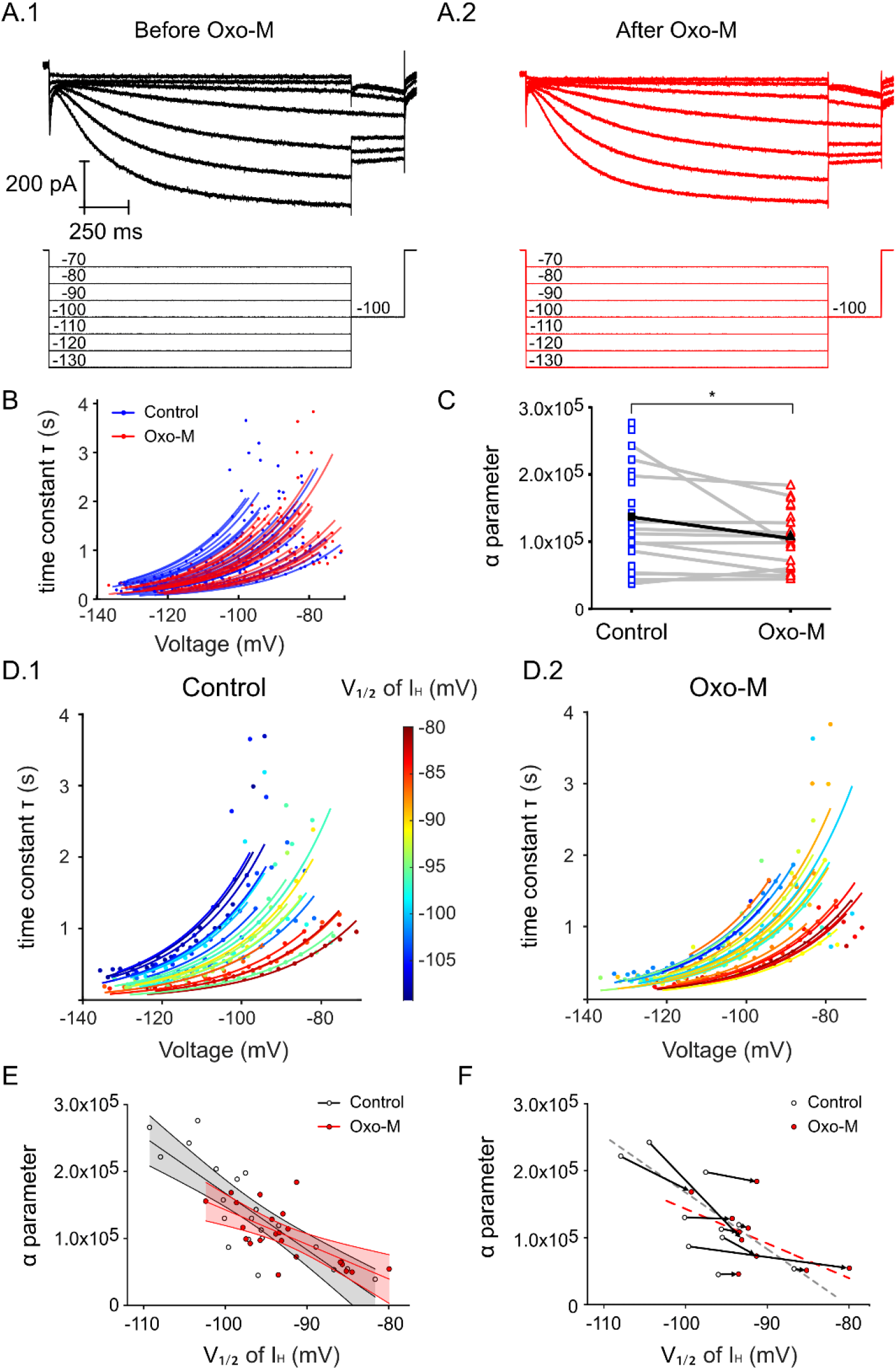
mAChR agonists increase the rate of HCN channel activation. **A.** Currents activated from voltage-steps from -130 to -70 mV in a single cell in control- solution (A.1, black) and in Oxo-M (B.2, red). **B.** the time constant τ derived from fits to whole cell currents is plotted as a function of the command voltage-step (V). A single-parameter exponential described in Eq. 2 of the methods was used to fit the τ(V) for each cell. τ(V) recorded in the control condition are shown in blue and cells from the Oxo-M condition are shown in in red. **C.** α from equation 2 is plotted for each cell before (blue square) and after Oxo-M (red triangle). Thin lines connect individual cells. The bold line indicates the average change in α. Oxo-M increased the activation rate of *I*_H_ as indicated y the reduction in α (* p=0.0309). **D.1.** *τ*(*V*) in control solution. **D.2.** *τ*(*V*) in Oxo-M. Points and lines are color-coded according to *V*_1/2_, with the most depolarized *V*_1/2_ values in red and the most hyperpolarized V_1/2_ values in blue. **E.** α values plotted against the *V*_1/2_ of each cell on the x-axis in control (black) and Oxo-M (red) conditions. Straight lines represent linear regression and 95% confidence interval estimates of control and Oxo-M groups in black and red, respectively. **F.** α and *V*_1/2_ measurements pairs from individual cells tested before (black) and after (red) Oxo-M administration. Arrows are drawn between two data points in each cell. Linear regressions in from 2E are shown in dotted lines in 2F.

We next considered whether the linear relationship between HCN channel activation rate and range remains intact after Oxo-M depolarizes the voltage-activation range of *I*_H_. Recall that in Figure 1F we described the correlation between *V*_1/2_ and α in the control condition; cells with more depolarized *V*_1/2_ also had faster activation kinetics. In Figures 3D.1 and 3D.2, we replot *τ*(*V*) in control and Oxo-M conditions with individual curves colored according to *V*_1/2_ (blue to red as *V*_1/2_ becomes more depolarized). The depolarizing shift in *V*_1/2_ by Oxo-M is evident in Figure D.2 by the decrease in number of cells with hyperpolarized *V*_1/2 -_in blue and an increase in the number of cells with depolarized *V*_1/2_ in red. The overall diversity in *V*_1/2_ values was unaffected (F(1,42)=0.0006, p=0.9802, Levene’s test, N=44). However, the activation rate is less diverse after Oxo-M, which is reflected in the significant reduction in variability in the activation-rate parameter *α*(F(1,42)=7.1294, p=0.0107, Levene’s Test, N=44). These results show that the effect of Oxo-M was strongest in cells with slower *I*_H_ kinetics, while cells with HCN channels that activate more quickly were less affected. While this may indicate that the effect of Oxo-M on the HCN channel activation rate is saturated in cells with faster baseline activation rates, our results suggest that the muscarinic signaling cascades affect the activation rate in some cells differently than others.

To assess how activating mAChR influences HCN channel kinetics in the broad population of cells recorded, we next directly compared the correlation between *I*_H_ voltage-activation range and activation rate before and after Oxo-M. As described in Fig. 1F, parameter α derived from the exponential fits to the voltage-activation range measured by *V*_1/2_ quantifies this dependence. Figure 3E adds the linear correlation between α and *V*_1/2_ in the Oxo-M condition. α is significantly dependent on *V*_1/2_ in both the control and Oxo-M conditions (r_control_(20)=-0.84, p<.0001,N=22, r_Oxo-M_(20)=-0.68, p=0.0005, n=22). However, the strength of the dependence between α and *V*_1/2_ is weaker in Oxo-M, as indicated by the shallower slope of the linear regression. This means that the *I*_H_ activation kinetics in relatively slow cells may have been more affected than the kinetics in cells that are relatively fast. This is particularly evident in individual cells where *I*_H_ was characterized in both control and Oxo-M conditions.

Figure 3F shows α in 10 cells that saw at least a 1 mV shift in *V*_1/2_. We found that only 2 cells saw a significant shift in both the activation range and speed of I_H_, while 8 cells reported shifts in the activation range only. Of the two cells that saw a large reduction in the activation rate, both had voltage activation ranges that were relatively hyperpolarized prior to mAChR activation (negative to -100 mV). These results show that the effect of muscarinic signaling cascades on HCN channels in VGN may vary depending on endogenous differences between cells such as cAMP concentration or HCN channel isoform, although identifying the mechanism is beyond the current scope of this article. In the following sections, we consider the extent to which HCN channel sensitivity to muscarinic signaling cascades is correlated with two features that are known to be heterogenous in VGN: depolarization-evoked firing patterns and cell size.

### Vestibular ganglion neurons that differ in firing pattern and cell size also have different HCN channel properties

The goal of the following analysis was test if HCN channel*s* properties and the HCN current’s sensitivity to mAChR activation varied by VGN sub-groups. Two criteria were used to classify VGN sub-groups*: 1)* cell size (as inferred from capacitance measurements) and 2) firing patterns in response to current injection. We considered whether *I*_H_ properties may vary according to cell size because of the following: 1) Afferents with larger cell bodies and axon diameter are more common in the central region of vestibular epithelia (Lysakowski and Goldberg, 1997), 2) afferents that project from the central regions are more likely to synapse with Type I hair cells with HCN channels that mediate to non-quantal transmission (Leonard and Kevetter, 2002; Contini et al., 2020) and 3) the HCN2 channel subunit, which is more sensitive to cAMP than other HCN channel subunits, is selectively expressed in a subset of large VGN (Almanza et al., 2012). We also assessed HCN channels in relation to firing patterns evoked in current-clamp mode. VGN with transient firing patterns are linked to irregular afferents from the central region of the vestibular sensory epithelia due to their ability to generate irregular firing patterns in response to pseudosynaptic stimuli (Kalluri et al., 2010; Ventura and Kalluri, 2019). Given that HCN channels may play different roles in afferents from the central and peripheral regions, these transients may also differ in their HCN channel properties.

### Defining vestibular ganglion neuron sub-groups based on firing patterns and cell size

VGN exhibit diverse responses to simple injections of depolarizing currents. Previous work has established that this diversity is driven by varied expression of low voltage_-_activated potassium channels (Iwasaki et al., 2008; Kalluri et al., 2010; Yoshimoto et al., 2015; Hight and Kalluri, 2016). In response to depolarizing current injection, VGN broadly show two distinct firing patterns: transient-firing or sustained-firing (Figure 4A). We classified VGN as either transient-firing (N=40) or sustained-firing (N=41) according to criteria described in Kalluri et al., 2010 and Iwasaki et al., 2008 (Iwasaki et al., 2008; Kalluri et al., 2010). Transient-firing VGN produce a single action potential in response to 200-ms current steps that reach or exceed spike threshold (Figure 4A.1, N=40). The responses of sustained-firing VGN are more variable and can be divided into three subgroups. Sustained-A neurons fire continuously throughout the current step (Figure 4B.1, N=5). Sustained-B neurons fire multiple action potentials (up to 9 in this study) that increase in frequency at larger current steps (Figure 4C.1, n=26). Sustained-C neurons fire a single action potential followed by voltage oscillations (Figure 4D.1 N=10). In contrast to positive current steps, large negative current steps produced a sharp hyperpolarization followed by a slow depolarization in all VGN. The resulting voltage ‘sag’ is indicative of the activation of net inward currents through HCN channels (see arrow in Figure 4A.1 through 4D.1). Figure 4E.1 shows the number of cells recorded with each firing pattern. The largest firing pattern group in the age range of this study had Transient firing patterns (N=40), while cells with Sustained-A firing patterns were rarely encountered (N=5).

**Figure 4.**
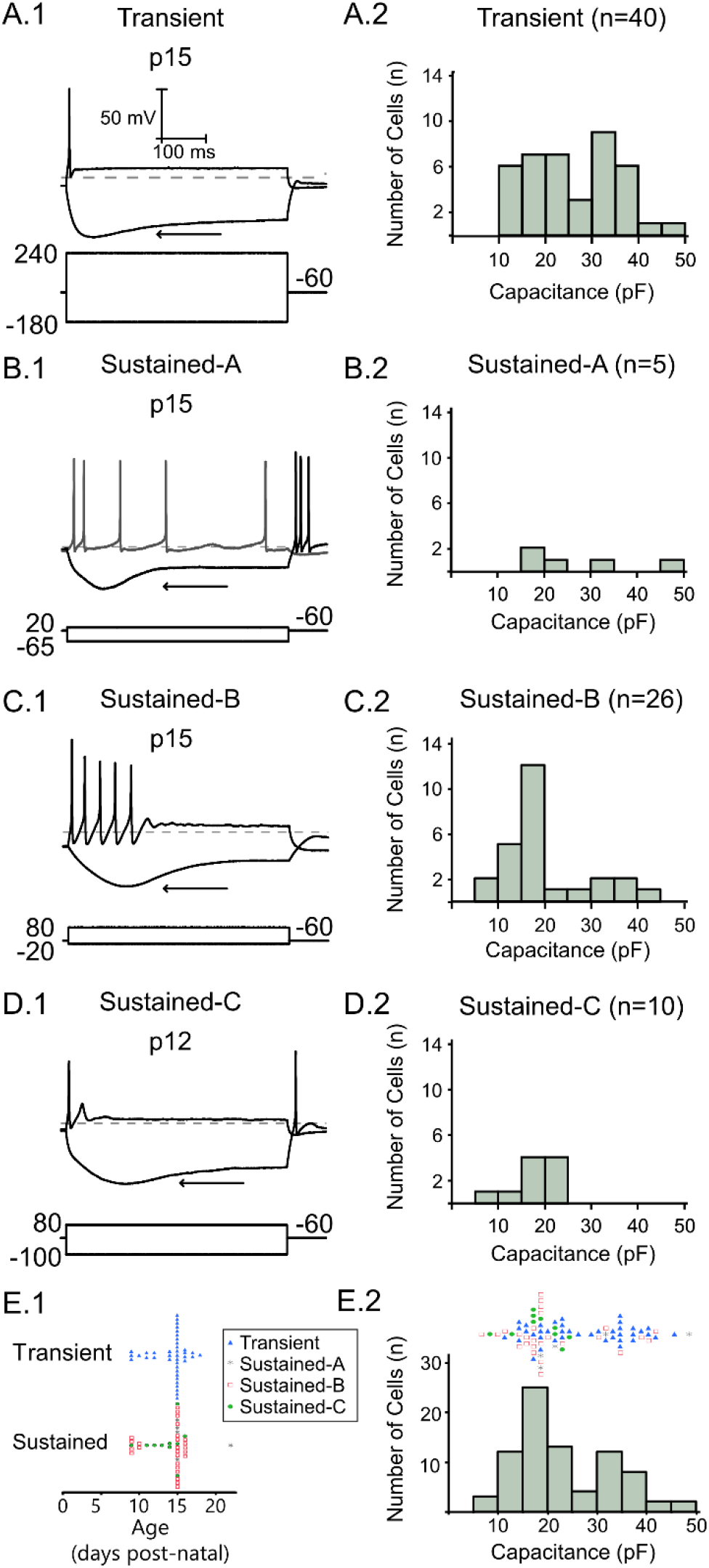
Vestibular ganglion neurons are heterogenous with different firing patterns in response to depolarizing current steps. Large hyperpolarizing current steps produce a voltage sag (arrow) driven by the hyperpolarization-activated mixed cationic current (*I*_H_). Scale bars in are consistent throughout all traces. Depolarizing current steps are the closest 20 pA current step to threshold in each trace. **A.1** Transient-firing neurons fire a single action potential at the onset of suprathreshold depolarizing current steps. The amplitude of the positive current step necessary to reach threshold is shown below each trace and the dashed line indicates -60 mV. **A.2.** Size was measured from membrane capacitance (*C*_m_). The size distribution of transient-firing neurons is bimodal with approximately 1/3 of all transients classified as large (> 30 pF). **B.1.** Sustained-A neurons fire continuously throughout the current step. **B.2.** Sustained-A firing neurons are mostly smaller than transient-firing VGN with 3 of the 5 cells recorded with a *C*_m_ less than 30 pF. **C.1** Sustained-B neurons fire multiple action potentials after the onset of the depolarizing current steps and are rapidly-adapting. **C.2.** Cells with sustained-B firing patterns varied from large to small but were no average smaller than transient-firing neurons. **D.1.** Sustained-C neurons fire a single action potential followed by voltage oscillations. **D.2.** All sustained-C neurons were in the - small size group. **E.1.** firing patterns in all cells (n=81) plotted against age in post-natal days. Transients are shown as blue triangles (n=40), sustained-A as gray asterisks (n=5), sustained-B as red squares (n=26), and sustained-C as green circles. **E.2.** Firing pattern of all 81 cells according to their size as measured by *C*_m_.

In disassociated VGN, membrane capacitance is an indirect measure of somatic size (Limón et al., 2005). We measured capacitance (*C*_m_) by first fitting a single exponential to the transient current evoked by a 5-mV depolarizing step and measuring the membrane time constant (*τ*_m_). *C*_m_ was calculated by dividing *τ*_m_ by the series resistance. On-line calculation of *C*_m_ was verified off-line using the average of multiple depolarizing steps recorded with whole-cell compensation turned off. Consistent with other studies, transient-firing VGN had higher capacitance than sustained-firing VGN (Kalluri et al., 2010; Ventura and Kalluri, 2019 p.20). Figures 4A.2 through 4D.2 show the distribution of cell size in VGN with different firing patterns. Figure 4E.2 shows that the distribution of capacitance values in 65 recorded cells. The overall distribution is bimodal with most cells having a C_m_ of 15 to 20 pF and another group of large cells between C_m_ between 25 and 30 pF C_m_. All large cells were classified as transient except for one large cell with a sustained-A firing pattern. In the following sections, we will classify VGN with capacitance values 30 pF or greater as large cells and those with Cm < 30 pF as small, which is the same size classification criteria used in other studies (Limón et al., 2005; Mercado et al., 2006; Almanza et al., 2012).

### The dependence of I_H_ activation range and kinetics on firing pattern

Figure 5A shows three representative examples of *I*_H_ current responses in VGN with transient, sustained-A, sustained-B and sustained-C firing patterns. We recorded whole-cell recording of I_H_ properties in only one cell with a sustained-A firing pattern as they are rarely found in this study’s age range, which spans the second and third post- natal weeks (Kalluri et al., 2010; Ventura and Kalluri, 2019). The *I*_H_ current densities of in all firing pattern groups were not statistically different (Current density transient = 12.2 ± 1.0 pA/pF, N = 29, sustained-A = 12.1 pA/pF, N=1, sustained-B = 12.0 ± 1.4, N=11, sustained-C = 11.0 ± 3.2 pA/pF, N=7, F(3,47)=0.0850, p=0.9679). However, larger cells had greater maximum currents to hyperpolarization (*I*_max_). Cells with transient-firing patterns had greater membrane capacitance (24.1 ± 1.8 pF) and *I*_max_ (285.8 ± 30.0 pA, N=29). VGN with sustained-B firing patterns were similarly larger than cells with sustained-C firing patterns (C_m_ Sustained-B = 20.1 ± 3.2 pF, N=11, C_m_ Sustained-C = 16.6 ± 2.0 pF, N=7) and had *I*_max_ (*I*_max_ Sustained-B = 256.2 ± 64.2 pF, N=11, *I*_max_ Sustained-C = 156.3 ± 22.9 pF). The only cell with a sustained-A firing pattern that we have *I*_H_ data on was between transient firing VGN and sustained-B firing VGN in terms of cell size and *I*_max_ (18.5 pF, 224.3).

**Figure 5.**
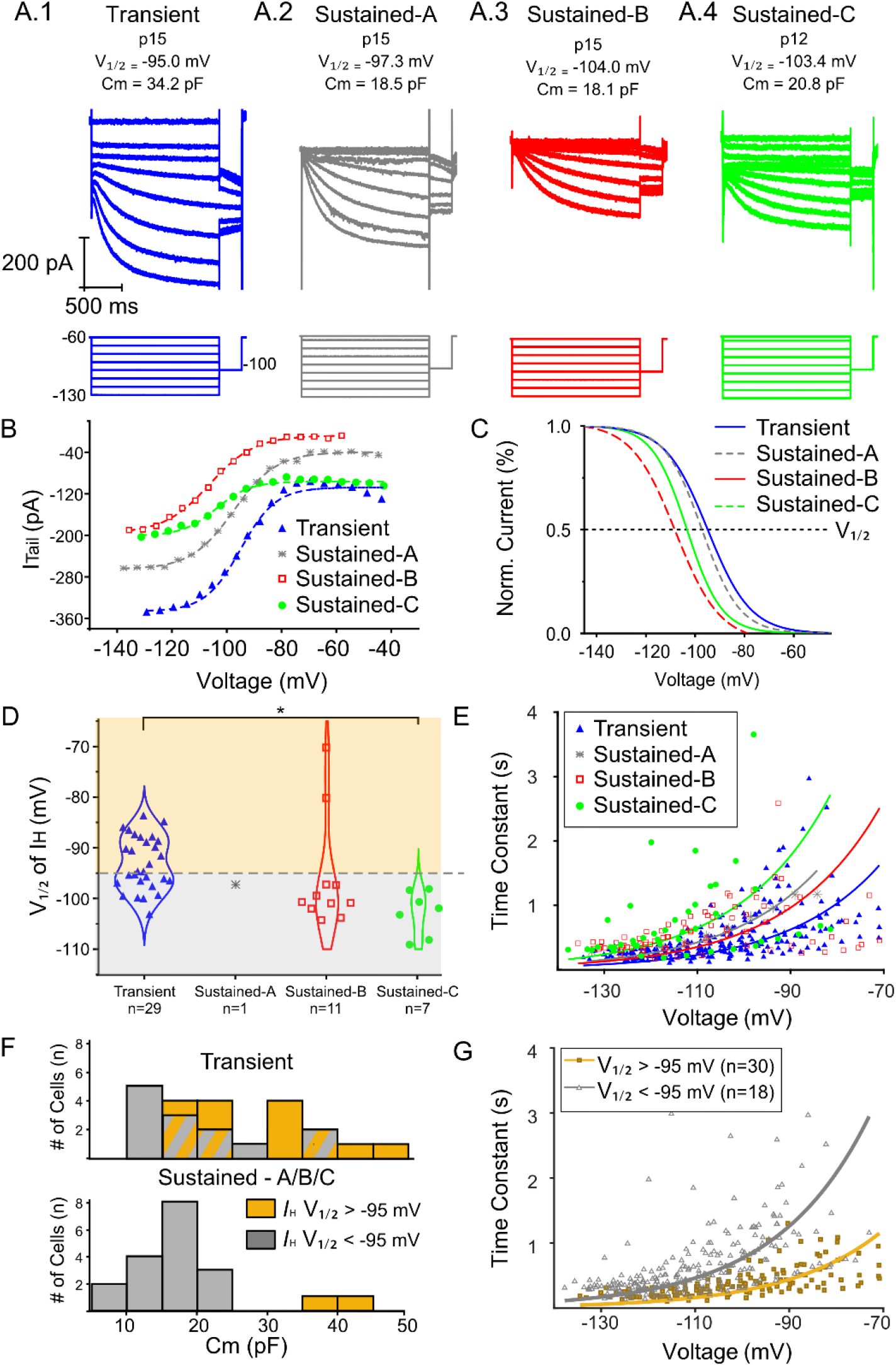
Heterogeneity of *I*_H_ activation properties as a function of firing pattern and cell size. **A.** Currents activated from voltage-steps from -135 to -60 mV followed by a -100 mV tail step from four representative different neurons with different firing patterns. Vertical gray line indicates where tail currents are measured. B-C and E show data from the individual neurons shown in A. Large transient-firing is shown in A.1, sustained-A in A.2, sustained-B in A.3, and sustained-C in A.4. Note that the sustained-A example shown in A.2. is the only cell in this group. In same neurons, **B.** tail-currents and **C.** *I*_H_ activation curves fit by Boltzmann function and normalized to the maximum conductance of *I*_H_ are shown. **D.** Violin-plots and individual data points on a normal distribution of *V*_1/2_ of *I*_H_ in each firing-pattern group are shown. *V*_1/2_ is more depolarized in transient-firing neurons than sustained-C neurons (* p=0.0084, Tukey’s HSD). Cells in which the *V*_1/2_ of *I*_H_ was positive of -95 mV were designated as HCN-enhanced (gold area of graph) and cells with *I*_H_ activation ranges more negative than - 95 mV were grouped as HCN-inactive (grey area of graph) **E.** A single-exponential was fit to each pre-conditioning step from -135 to -70 mV from cells in each firing pattern and the time constant τ from each step is plotted as a function of voltage. Cells with transient firing patterns are shown in blue triangles, sustained-A in gray asterisks, sustained-B in red squares and sustained-C in green circles. **F.** C_m_ distribution in transient-firing (top) and sustained-A, B, and C firing (bottom) cells. The number of cells classified as HCN-enhanced is shown in the histogram in gold while cells defined as HCN-inactive are shown in gray. Overlap between HCN-enhanced and HCN-inactive in the distribution of *C*_m_ measurement is shown as grey and gold stripes. **G**. *τ* constant in all cells are shown as either HCN-enhanced (gold squares, n=18) or HCN-inactive (gray triangles, n=30).

We next tested whether cells with different firing patterns have different *I*_H_ voltage-activation ranges. In the example traces, the cell with the transient firing pattern has a more depolarized *I*_H_ voltage-activation range than the cells with sustained-A, sustained-B or sustained-C firing patterns (Figure 5C). These examples are representative of the relationship between firing pattern and *I*_H_ voltage-activation range with transient-firing cells having more depolarized *I*_H_ voltage-activation range (transient V_1/2_ = -93.3 ± 1.0 mV, n = 29) than cells with sustained-C firing patterns (-103.0 ± 1.7, n=7, firing pattern effect on V_1/2_ : ANOVA F(3,44)=3.86, p=0.015, ANOVA, Difference in V_1/2_ between transient and sustained-C VGN: Tukey’s HSD, p=0.0084). However, only about half of transient-firing VGN had *I*_H_ voltage-activation ranges that were more depolarized than cells with sustained-B or sustained-C firing patterns (Figure 5D).

Consistent with their more hyperpolarized *V*_1/2_, *I*_H_ activated slower is sustained-C firing neurons than in transient-firing neurons, although the latter group had considerable variability. Two cells with the most depolarized *I*_H_ voltage-activation range and fastest kinetics in the study had sustained-B firing patterns.

We classified any cell with an *I*_H_ activation range more positive than -95 mV as an HCN-enhanced cell (Figure 5D, above horizontal dotted line). In Figure 5F, we plotted the distribution of cell size. Different colored bars indicate the number of cells with HCN-enhanced (yellow) or HCN-inactive (gray). HCN-enhanced cells were larger on average than HCN-inactive cells regardless of firing pattern (C_m_ HCN-enhanced = 29.5 ± 2.3 pF, n=18; C_m_ HCN-inactive 17.5 ± 1.1 pF, N=30; t(47)=27.9, p<.0001). The two HCN-enhanced, sustained-B firing VGN were significantly larger than all other sustained-A, B and C-firing VGN in our sample. These results show that VGN large cell size and transient firing patterns are two related features that correlate with *I*_H_ voltage- activation range. One possibility is that these large cells express HCN channel subunits with greater sensitivity to cAMP as suggested in the preferential expression of the HCN2 subunit in large cell bodies of the vestibular epithelia (Almanza et al., 2012).

We also tested whether cells with a depolarized *I*_H_ activation range (HCN- enhanced) have faster kinetics than cells in which the *I*_H_ activation range is hyperpolarized (HCN-inactive). Note that the HCN-enhanced and HCN-inactive groups contain a mix of both transient and sustained-firing cells. The time course of I_H_ was measured by fitting single-exponential curves to the I_H_ currents elicited by voltage steps between -130 mV and -70 mV. In Figure 5F, we plotted the exponent τ as a function of voltage in HCN-enhanced cells (yellow closed squares) and HCN-inactive cells (open gray triangles). The voltage-dependent relationship between the time course of *I*_H_ activation in HCN-enhanced and HCN-inactive cells were fit using an exponential equation with a single parameter (Equation 2, See Materials and Methods). HCN- enhanced VGN had faster *I*_H_ kinetics than HCN-inactive VGN. Together, these results underline the diversity in HCN channel properties and suggest that this diversity is related to cell size and firing pattern. Given that perforated patch revealed this diversity suggests that this technique may preserve intracellular factors such as cAMP that depolarize the activation range of HCN channels in some VGN. These HCN-enhanced VGN were mostly large and transient-firing, although small transient-firing and large sustained-firing VGN were also included in this group.

### Muscarinic modulation of I_H_ shapes firing pattern in VGN with sustained-firing patterns

Given that VGN with different firing patterns and size also differ in their voltage-dependent properties of *I*_H_ at baseline, an important question is whether these differences shape the sensitivity to mAChR activation. We first assessed the sensitivity of firing pattern to mAChR activation in 6 cells that were tested in all three bath solutions: control, linopirdine, and linopirdine plus Oxo-M. We selected three example cells in which we saw differing degrees of responsiveness to Oxo-M. In each cell we examined changes in the voltage activation range of *I*_H_, resting membrane potential (RMP) and step-evoked firing patterns. Figure 6A.1 shows an example of a large, transient-firing cells in all three conditions. Although Oxo-M shifted the *I*_H_ voltage activation range by approximately 5 mV, the firing pattern was virtually unchanged after adding linopirdine and after adding Oxo-M. The absence of an *I*_H_-mediated effect on transient firing patterns is consistent with previous studies showing that in some neurons transient-firing patterns are dominated by Kv1 mediated low-voltage activated potassium channels that remain active after treatment with Oxo-M (Iwasaki et al., 2008; Kalluri et al., 2010; Hight and Kalluri, 2016). We have previously shown that these low-voltage activated potassium channels counteract the excitatory effects of HCN channel activation in a model cell (Ventura and Kalluri, 2019).

**Figure 6.**
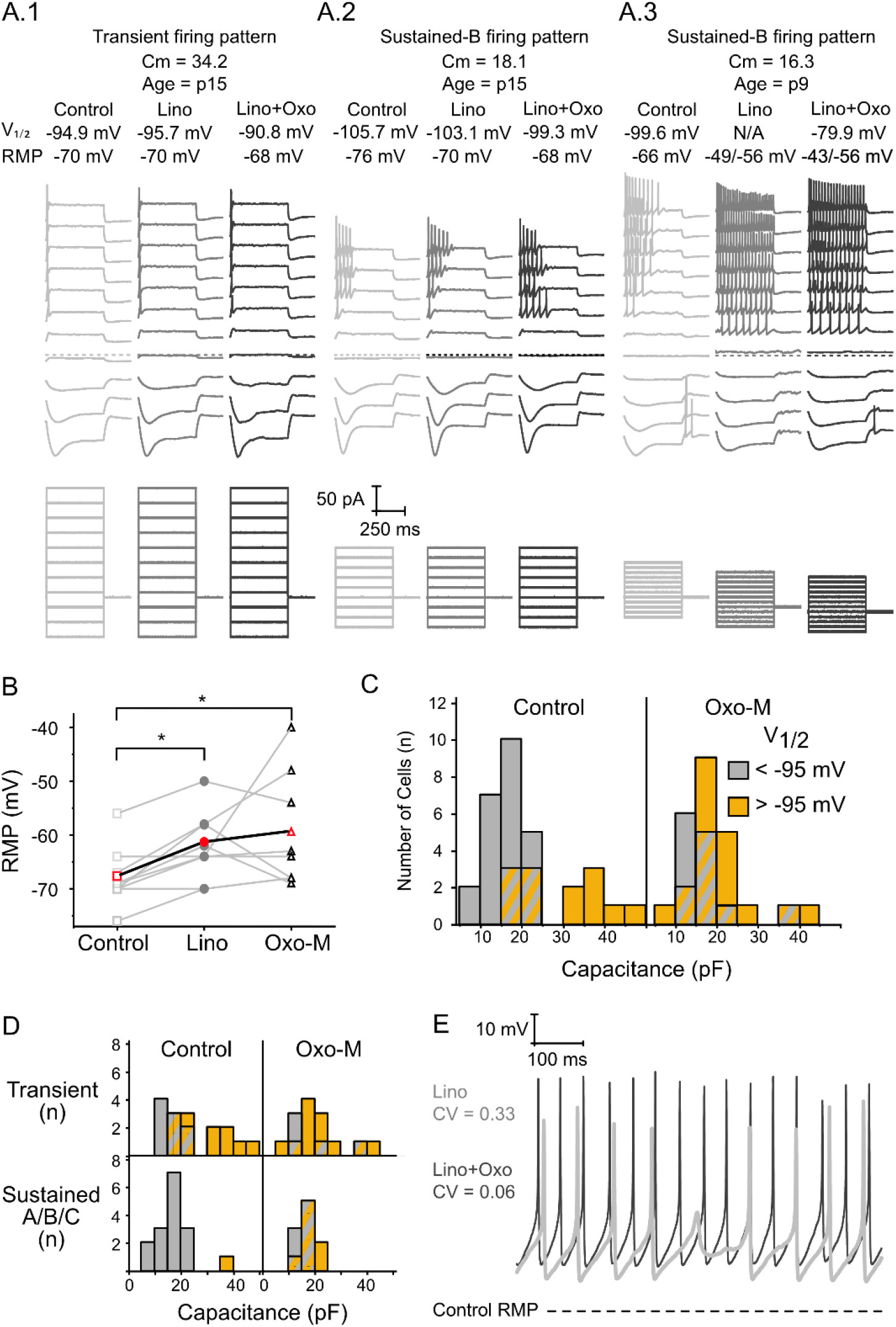
Heterogeneity in the impact of activation mAchR on firing patterns. **A.1**. Large transient-firing cell that saw no change in firing pattern despite a moderate positive shift in *I*_H_ activation range, **A.2**, Sustained-B with moderate effect, **A.3.** Sustained-B with large effect. Voltage response are shown in a stacked array separated by 10 mV. Dashed horizontal-line drawn at -60 mV in all three cells Traces are shown at baseline control (light grey), after linopirdine (dark grey) and after linopirdine and oxo-M (black). Note the cell in A.3. was held at -59 mV in both lino and lino+oxo conditions **B.** Resting membrane potential is plotted for each cell before (square), after linopirdine (circle), and after linopirdine and oxo-M (triangle). Thin lines connect individual cells. The bold line and red data points indicates the average change RMP in each condition. Both lino and lino+oxo-M significantly depolarized RMP relative to control (p>.05, Tukey HSD). **C.** The voltage-activation ranges of I_H_ in small, sustained-firing VGN were most sensitive to mAChR activation. C_m_ distribution in control (left) and Oxo-M conditions. Bars are color coded for *I*_H_ *V*_1/2_ values more positive (HCN-enhanced, gold) or negative (HCN-inactive, gray) than -95 mV. Overlap of HCN-enhanced and HCN-inactive distributions are shown as gold and gray stripes. Note the increase in yellow on the distribution of small cells in the Oxo-M condition. **D.** C_m_ distribution in transient-firing (top) and sustained-A, B or C-firing (below) cells in control (left) and oxo-M (right) conditions. Note that only 1 sustained-firing cell is classified as HCN-enhanced in the control condition, while approximately of the sustained-firing VGN are classified as control after Oxo-M. **E**. Spontaneous-firing in the cell shown in A.3 a in the presence of linopirdine alone (Gray thick trace) and in the presence of linopridine and Oxo-M (Black thin trace). Note the reduction in spike failure and the increase in the regularity of spikes after Oxo-M is added to the bath. Cell was not spontaneously active in the control condition. RMP in control condition is shown as a dotted horizontal line. This increase in regularity is reflected in the decrease in coefficient of variability (CV) from 0.33 after Lino to 0.06 after Lino+Oxo, which is similar to regular firing observed *in vivo*.

In cells with sustained-B firing patterns, *I*_KL_ is likely comprised in part by KCNQ channels that are closed by Oxo-M and Lino (Kalluri et al., 2010). In this case, the lack of inhibition by *I*_KL_ may allow HCN channels to contribute to VGN firing patterns. Figure 6A.2 shows a Sustained-B with an *I*_H_ voltage activation range that also shifted in the depolarizing direction by 3-5 mV, but in this case the cell fired 5 spikes near threshold after Oxo-M while only a single action potential was fired in control and lino conditions.

In some neurons, the resting potential of the neuron depolarized significantly in the presence of Oxo-M. The cell shown in Figure 6A.3 saw the largest shift in *I*_H_ voltage-activation range after adding Oxo-M at ∼11 mV. The depolarization from the closure of KCNQ was so significant in this cell (from -66 mV to -49 mV) such that it was unable to fire an action potential after the addition of linopirdine. The addition of Oxo-M to the bath solution further depolarized the membrane potential (-43 mV). The inability to fire an action potential presumably occurred due to sodium channel inactivation at this membrane potential since the cell fired successfully when held at -56 mV in both lino and oxo-M conditions (dark gray and black traces, Figure 6A.3). When held at a constant -56 mV, the cell fired slightly fewer spikes near threshold (7 spikes after lino when compared to 6 spikes after Oxo-M). In this case, the depolarization of resting membrane potential and the resulting sodium channel activation may have obscured changes in firing pattern. In Figure 6B, we plotted the resting membrane potential in 8 cells as they transitioned between bath solutions with control, linopirdine, and linopirdine and Oxo-M. Linopirdine and Oxo-M significantly depolarized RMP (F(2,14)=7.51, p=0.0061) as the bath solution was switched; (RMP_Control_ = -68.3 ± 1.9; after linopirdine RMP_Lino_ = -59.9 ± 3.2 mV, p > .05, Tukey HSD) and then again after linopirdine and Oxo-M (RMP_Lino+Oxo-M_ = -57.8 ± 4.4 mV, p > .05, Tukey HSD).

Next, we compared the relationship between cell size and *I*_H_ half-activation properties in cells from both control and Oxo-M conditions (Figure 6C). In the control condition, cells with depolarized *I*_H_ properties (V_1/2_ > - 95) tend to be larger cells. In contrast, after Oxo-M there is no obvious relationship between cell size and *I*_H_ voltage- activation properties since the number of cells with depolarized *I*_H_ properties is evenly distributed among different sized cells. This suggests that the cells that are most-sensitive to Oxo-M are the smaller cells. Given the relationship between cell size and firing pattern, we next compared the activation properties of cells that received Oxo-M or control within each firing pattern group (transient or sustained-A/B/C). Figure 6D shows that the effect of Oxo-M on transient-firing VGN was less dramatic, as cells in both control (V_1/2_: -93.5 ± 1.3 mV, n=21) and oxo-M conditions (V_1/2_: -92.7 ± 1.2 mV, n=17) were almost equally depolarized. In contrast, VGN with sustained-A/B/C firing patterns, which also tend to be small cells, saw the largest differences between control (-100.9 ± 1.1 mV, n=7) and Oxo-M conditions (-93.3 ± 1.7, n=9). Together, these results suggest that the sensitivity of *I*_H_ to muscarinic signaling cascades is dependent on both cell size and firing pattern. This suggests that there is considerable heterogeneity in either the intracellular factors and/or in HCN channel sub-units that determine *I*_H_ properties at baseline. For example, transient-firing VGN have depolarized *I*_H_ properties at baseline and may be saturated prior to any mAChR activation-induced shift in V_1/2_. In contrast, the *I*_H_ activation range in sustained-firing cells is hyperpolarized with room to be depolarized via intracellular signaling pathways.

The powerful impact of activating HCN on top of closing KCNQ was apparent in a handful of cells whose firing patterns were affected by Oxo-M in a way that was not observed following linopirdine. The most striking example is shown in Figure 6E. A sustained-B neuron in control conditions became spontaneously active after we applied linopirdine (Figure 6E, gray). Though spontaneously active, the firing pattern showed some burstiness as some spikes failed to fully form. The firing pattern remained spontaneous when Oxo-M was added (Figure 6E, black). However, notice that there were fewer failed spikes producing a striking increase in the regularity inter-spike intervals (CV_Lino_ = 0.33, CV_Lino+Oxo_ = 0.06). In this case, Oxo-M induced the most regular firing we’ve recorded *in vitro* and the CV value is comparable to the most regular afferent firing patterns observed *in vivo* (Goldberg et al., 1990, Ventura and Kalluri, 2019). Therefore, the muscarinic receptor agonist, Oxo-M, boosted the regularity of spiking to beyond that which resulted from simply closing KCNQ. This is likely the result of enhancing *I*_H_ due to the depolarization of HCN channel activation. This role of *I*_H_ is akin to its famous role in promoting pace-making in other cells, including the heart (Harvey and Belevych, 2003; DiFrancesco Dario, 2010).

## Discussion

### An endogenous mechanism for depolarizing HCN channels

HCN channels are thought to contribute to vestibular afferent signaling by promoting non-quantal transmission and regular spike-timing (Horwitz et al., 2014; Contini et al., 2020). However, previous in vitro characterizations from isolated ganglion neurons indicated that the HCN current, *I*_H_, is mostly unavailable for shaping vestibular neuron activity (Hight and Kalluri, 2016) due to its hyperpolarized voltage activation range (Chabbert et al., 2001; Almanza et al., 2012). In vitro experiments also show that the activation range of HCN channels in vestibular afferents can be depolarized into a physiologically useful range by artificially elevating the concentration of intracellular second messengers like cAMP (e.g., Horwitz et al., 2014; Ventura and Kalluri 2019). Such modulation of HCN can dramatically alter patterns of afferent activity in immature neurons (Horwitz et al., 2014). Our results show cholinergic efferent inputs drive a mechanism for naturally activating HCN channels. This would make more *I*_H_ current available for shaping afferent activity. Given recent evidence that the cholinergic vestibular efferents are tonically active (Raghu et al., 2019), *I*_H_ may be more available in vivo than indicated by previous in vitro work. In summary, our results show that HCN channels are targeted by the same muscarinic signaling cascades that are already known to influence the firing rate of vestibular afferent neurons by closing KCNQ channels (Pérez et al., 2009; Holt et al., 2017).

G-protein coupled signaling cascades triggered by mAChR receptors affect the availability of cAMP and PIP_2_ and accordingly the *I*_H_ voltage-activation range (Robinson and Siegelbaum, 2003; Pian et al., 2007). Whether mAChR activation upregulates intracellular cAMP or PIP_2_ depends on the receptor subtype, all five of which (M1 through M5) are expressed in vestibular neurons (Li et al., 2007). Muscarinic acetylcholine receptors that contain M1, M3 and M5 subtypes couple to the Gαq/11 G-protein and activate the phospholipid, phosphatidylinositol-4,5-bisphosphate (PIP_2_) (Brown, 2010). Activation of M1-receptors in expression systems depolarizes the *I*_H_ activation range through a PIP_2_-related mechanism (Pian et al., 2007). The M2 and M4 subtypes release Gi/Go G-proteins that can either inhibit or stimulate cAMP activity. Increasing cAMP concentration in VGN shifts the *I*_H_ activation range in the positive direction (Almanza et al., 2012; Ventura and Kalluri, 2019). Whether efferent modulation of HCN channels is mediated by cAMP or PIP_2_ is unclear, but multiple signaling pathways may be involved because VGN express diverse mAChR subtypes.

### I_H_ activation range in VGN is cell size and firing pattern-dependent

VGN are remarkably diverse in their cell size, discharge pattern and ion-channel composition (Iwasaki et al., 2008; Kalluri et al., 2010; Almanza et al., 2012; Yoshimoto et al., 2015; Hight and Kalluri, 2016; Ventura and Kalluri, 2019). Our results provide novel insight into how these categories overlap with HCN channel properties, which we found to be more variable than previously characterized. The half-activation voltage of *I*_H_ ranged from -70 mV to -109 mV. Previous studies did not find *I*_H_ activation ranges near -70 mV in VGN (Chabbert et al., 2001; Almanza et al., 2012; Yoshimoto et al., 2015; Ventura and Kalluri, 2019). The likely difference between this and previous studies is that we used perforated-patch methods to characterize *I*_H_ (8 cells across two previous studies compared to 48 cells in this study used perforated-patch; (Almanza et al., 2012; Ventura and Kalluri, 2019). Rupture-patch techniques used in previous studies cause the intracellular composition to dialyze with the contents of the recording pipette. The resulting time-dependent rundown in second messengers likely hyperpolarizes the activation range of HCN and reduces the cell-to-cell variability in HCN channel properties observed here.

### Functional Significance

Efferent control of *I*_H_ is significant because of the relevance of HCN channels to two hallmarks features of vestibular afferent signaling, non-quantal transmission and spike-timing regularity. HCN channels are believed to be essential for producing a form of fast (non-quantal) transmission between vestibular hair cells and the calyceal synapses of vestibular afferents (Contini et al., 2020). This ultrafast inward current is thought to facilitate the rapid transmission of sensory information from hair cells to afferent neurons (Gersdorff et al., 2020). Our results show that large VGN with transient firing patterns, which likely include the somata of pure calyx afferents from the central zones of vestibular epithelia (Kevetter and Leonard, 2012), have HCN channels that are naturally more depolarized than HCN channels in small ganglion neurons. The more depolarized HCN channels may therefore be important for supporting non-quantal transmission.

### Modulation of non-quantal transmission

Efferent control of HCN channels has the potential to shape non-quantal transmission in vestibular afferents. HCN channels provide the pathway for resistive coupling between Type I hair cells and post-synaptic calyces (Yamashita and Ohmori, 1991; Songer and Eatock, 2013; Highstein et al., 2014; Contini et al., 2020). Resistive coupling results when K+ released from type I hair cells accumulates in the narrow synaptic cleft of the calyx and flows into the afferent through HCN channels (Goldberg, 1996; Contini et al., 2020). The size of this current is proportional to the resting HCN conductance on the inner face of the afferent. Efferent terminals synapse with the outer face of the calyx (Lysakowski and Goldberg, 1997), which express muscarinic M1, M2, M4 and M5 receptors (Li et al., 2007). To potentiate non-quantal transmission, mAChRs on the outer face would have to shift the activation range of HCN channels on the inner face of the calyx. Although it remains to be empirically tested, it is plausible for the intracellular signaling cascades triggered by mAchR receptor activation to impact remotely located HCN channels.

### Dual impact of mAchR on KCNQ and HCN channels is necessary for promoting regularity

The depolarizing currents of HCN channels are believed to be instrumental toward generating the strikingly regular inter-spike intervals found in many vestibular afferents (Horwitz et al., 2014; Yoshimoto et al., 2015). This is akin to HCN channels’ role in cardiac pacemaking (DiFrancesco and Tromba, 1988). Evidence supporting this hypothesis is that highly regular-spike patterns become more prevalent with post-natal development as HCN channel expression grows and matures (Yoshimoto et al., 2015) and spike-timing regularity is sensitive to pharmacological manipulation of HCN channel activity in neonatal vestibular afferents (Horwitz et al., 2014).

Vestibular ganglion neurons containing low-voltage gated potassium currents produce irregular-timed spike patterns in response to simulated synaptic currents (Kalluri et al., 2010). Although HCN channels promote pace-making in other systems (Pape and McCormick, 1989; DiFrancesco, 1993), their impact on vestibular afferent spike timing likely depends on whether the afferent also expresses low-voltage activated potassium channels. HCN channels interact with low-voltage gated potassium currents in auditory brainstem neurons to enhance phasic-firing (Oertel et al., 2000; Rothman and Manis, 2003; McGinley and Oertel, 2006; Cao and Oertel, 2011; Khurana et al., 2012). Similarly, Ventura and Kalluri (2019) showed that HCN channels amplify transient firing in disassociated vestibular ganglion neurons by shifting the membrane potential towards the activation range of potassium channels (Ventura and Kalluri, 2019). Thus, HCN channels enhance transient-spiking and irregular-timed spiking in neurons that contain *I*_KL_, even when the HCN channel activation is depolarized by artificially increasing the intracellular concentration of cAMP (Ventura and Kalluri, 2019). In a model, Ventura and Kalluri (2019) demonstrated that HCN channels could only promote regular-timed firing in simulated neurons that don’t have *I*_KL_.

Most mature vestibular ganglion neurons are likely to have some *I*_KL_ due to the upregulation of KCNQ channels with maturation (Rocha-Sanchez et al., 2007). This suggests that a developmental upregulation of HCN channels alone is unlikely to account for an increase in the prevalence of regular spiking with maturation. However, since vestibular efferent neurons are believed to be tonically active (Sadeghi et al., 2009; Raghu et al., 2019), the simultaneous closure of KCNQ channels and enhancement of HCN channel activation via the muscarinic receptor cascade provides a plausible scenario for promoting highly-regular firing.

Our results suggest that spike-timing regularity may not be a fixed property for all vestibular neurons, but dynamically regulated by efferent inputs. Efferent activity may promote ion channel configurations that favor both regular and irregular spike-timing, depending on the balance between low-voltage gated potassium currents and the activation range of HCN currents. Simultaneously closing KCNQ and enhancing HCN channels would promote regular-timed spiking. This means that some neurons may be capable of firing at both regular and irregular intervals depending on the strength of the efferent input. Spike-timing regularity may be less flexible in other neurons if they contain low-voltage gated potassium channels (such as Kv1) that are not easily closed by activating mAChR signaling (Iwasaki et al., 2008; Kalluri et al., 2010). The closure of KCNQ and enhancement of *I*_H_ would increase firing rate, but Kv1 mediated currents would still drive transient-spiking in response to injected current steps and produce irregular firing patterns in response to synaptic drive (as predicted by modeling, Ventura and Kalluri, 2019). Thus, efferent-mediated modulation is likely to be more nuanced than previously appreciated because of its dual impact on two ion channel groups known to shape features of vestibular afferent excitability and timing.

## References

Almanza A, Luis E, Mercado F, Vega R, Soto E (2012) Molecular identity, ontogeny, and cAMP modulation of the hyperpolarization-activated current in vestibular ganglion neurons. J Neurophysiol 108:2264–2275.

Barry PH (1994) JPCalc, a software package for calculating liquid junction potential corrections in patch-clamp, intracellular, epithelial and bilayer measurements and for correcting junction potential measurements. J Neurosci Methods 51:107–116.

Biel M, Wahl-Schott C, Michalakis S, Zong X (2009) Hyperpolarization-Activated Cation Channels: From Genes to Function. Physiol Rev 89:847–885.

Brown DA (2010) Muscarinic Acetylcholine Receptors (mAChRs) in the Nervous System: Some Functions and Mechanisms. J Mol Neurosci 41:340–346.

Cao X-J, Oertel D (2011) The magnitudes of hyperpolarization-activated and low- voltage-activated potassium currents co-vary in neurons of the ventral cochlear nucleus. J Neurophysiol 106:630–640.

Chabbert C, Chambard JM, Sans A, Desmadryl G (2001) Three Types of Depolarization-Activated Potassium Currents in Acutely Isolated Mouse Vestibular Neurons. J Neurophysiol 85:1017–1026.

Contini D, Holstein GR, Art JJ (2020) Synaptic cleft microenvironment influences potassium permeation and synaptic transmission in hair cells surrounded by calyx afferents in the turtle. J Physiol 598:853–889.

DiFrancesco D (1993) Pacemaker mechanisms in cardiac tissue. Annu Rev Physiol 55:455–472.

DiFrancesco D, Tromba C (1988) Muscarinic control of the hyperpolarization-activated current (if) in rabbit sino-atrial node myocytes. J Physiol 405:493–510.

DiFrancesco Dario (2010) The Role of the Funny Current in Pacemaker Activity. Circ Res 106:434–446.

Eatock RA, Songer JE (2011) Vestibular hair cells and afferents: two channels for head motion signals. Annu Rev Neurosci 34:501–534.

Gersdorff H von, Iversen MM, Rabbitt RD (2020) Keeping your eye on the ball. J Physiol 598:623–624.

Goldberg JM (1996) Theoretical analysis of intercellular communication between the vestibular type I hair cell and its calyx ending. J Neurophysiol 76:1942–1957.

Harvey RD, Belevych AE (2003) Muscarinic regulation of cardiac ion channels. Br J Pharmacol 139:1074–1084.

Highstein SM, Holstein GR, Mann MA, Rabbitt RD (2014) Evidence that protons act as neurotransmitters at vestibular hair cell–calyx afferent synapses. Proc Natl Acad Sci 111:5421–5426.

Hight AE, Kalluri R (2016) A biophysical model examining the role of low-voltage- activated potassium currents in shaping the responses of vestibular ganglion neurons. J Neurophysiol 116:503–521.

Holt JC, Jordan PM, Lysakowski A, Shah A, Barsz K, Contini D (2017) Muscarinic Acetylcholine Receptors and M-Currents Underlie Efferent-Mediated Slow Excitation in Calyx-Bearing Vestibular Afferents. J Neurosci 37:1873–1887.

Horwitz GC, Risner-Janiczek JR, Holt JR (2014) Mechanotransduction and hyperpolarization-activated currents contribute to spontaneous activity in mouse vestibular ganglion neurons. J Gen Physiol 143:481–497.

Hughes S, Marsh SJ, Tinker A, Brown DA (2007) PIP2-dependent inhibition of M-type (Kv7.2/7.3) potassium channels: direct on-line assessment of PIP2 depletion by Gq-coupled receptors in single living neurons. Pflüg Arch - Eur J Physiol 455:115–124.

Iwasaki S, Chihara Y, Komuta Y, Ito K, Sahara Y (2008) Low-Voltage-Activated Potassium Channels Underlie the Regulation of Intrinsic Firing Properties of Rat Vestibular Ganglion Cells. J Neurophysiol 100:2192–2204.

Kalluri R, Xue J, Eatock RA (2010) Ion Channels Set Spike Timing Regularity of Mammalian Vestibular Afferent Neurons. J Neurophysiol 104:2034–2051.

Khurana S, Liu Z, Lewis AS, Rosa K, Chetkovich D, Golding NL (2012) An Essential Role for Modulation of Hyperpolarization-Activated Current in the Development of Binaural Temporal Precision. J Neurosci 32:2814–2823.

Leonard RB, Kevetter GA (2002) Molecular probes of the vestibular nerve: I. Peripheral termination patterns of calretinin, calbindin and peripherin containing fibers. Brain Res 928:8–17.

Li GQ, Kevetter GA, Leonard RB, Prusak DJ, Wood TG, Correia MJ (2007) Muscarinic acetylcholine receptor subtype expression in avian vestibular hair cells, nerve terminals and ganglion cells. Neuroscience 146:384–402.

Limón A, Pérez C, Vega R, Soto E (2005) Ca2+-Activated K+-Current Density Is Correlated With Soma Size in Rat Vestibular-Afferent Neurons in Culture. J Neurophysiol 94:3751–3761.

Lysakowski A, Goldberg JM (1997) A regional ultrastructural analysis of the cellular and synaptic architecture in the chinchilla cristae ampullares. J Comp Neurol 389:419–443.

McGinley MJ, Oertel D (2006) Rate thresholds determine the precision of temporal integration in principal cells of the ventral cochlear nucleus. Hear Res 216–217:52–63.

Mercado F, López IA, Acuna D, Vega R, Soto E (2006) Acid-Sensing Ionic Channels in the Rat Vestibular Endorgans and Ganglia. J Neurophysiol 96:1615–1624.

Oertel D, Bal R, Gardner SM, Smith PH, Joris PX (2000) Detection of synchrony in the activity of auditory nerve fibers by octopus cells of the mammalian cochlear nucleus. Proc Natl Acad Sci 97:11773–11779.

Pape H-C, McCormick DA (1989) Noradrenaline and serotonin selectively modulate thalamic burst firing by enhancing a hyperpolarization-activated cation current. Nature 340:715–718.

Pérez C, Limón A, Vega R, Soto E (2009) The muscarinic inhibition of the potassium M- current modulates the action-potential discharge in the vestibular primary-afferent neurons of the rat. Neuroscience 158:1662–1674.

Pian P, Bucchi A, DeCostanzo A, Robinson RB, Siegelbaum SA (2007) Modulation of cyclic nucleotide-regulated HCN channels by PIP2 and receptors coupled to phospholipase C. Pflüg Arch - Eur J Physiol 455:125–145.

Raghu V, Salvi R, Sadeghi SG (2019) Efferent Inputs Are Required for Normal Function of Vestibular Nerve Afferents. J Neurosci 39:6922–6935.

Robinson RB, Siegelbaum SA (2003) Hyperpolarization-Activated Cation Currents: From Molecules to Physiological Function. Annu Rev Physiol 65:453–480.

Rocha-Sanchez SMS, Morris KA, Kachar B, Nichols D, Fritzsch B, Beisel KW (2007) Developmental expression of Kcnq4 in vestibular neurons and neurosensory epithelia. Brain Res 1139:117–125.

Rothman JS, Manis PB (2003) The Roles Potassium Currents Play in Regulating the Electrical Activity of Ventral Cochlear Nucleus Neurons. J Neurophysiol 89:3097–3113.

Sadeghi SG, Chacron MJ, Taylor MC, Cullen KE (2007) Neural Variability, Detection Thresholds, and Information Transmission in the Vestibular System. J Neurosci 27:771–781.

Sadeghi SG, Goldberg JM, Minor LB, Cullen KE (2009) Efferent-Mediated Responses in Vestibular Nerve Afferents of the Alert Macaque. J Neurophysiol 101:988–1001.

Songer JE, Eatock RA (2013) Tuning and Timing in Mammalian Type I Hair Cells and Calyceal Synapses. J Neurosci 33:3706–3724.

Ventura CM, Kalluri R (2019) Enhanced Activation of HCN Channels Reduces Excitability and Spike-Timing Regularity in Maturing Vestibular Afferent Neurons. J Neurosci 39:2860–2876.

Wainger BJ, DeGennaro M, Santoro B, Siegelbaum SA, Tibbs GR (2001) Molecular mechanism of cAMP modulation of HCN pacemaker channels. Nature 411:805– 810.

Yamashita M, Ohmori H (1991) Synaptic bodies and vesicles in the calix type synapse of chicken semicircular canal ampullae. Neurosci Lett 129:43–46.

Yoshimoto R, Iwasaki S, Takago H, Nakajima T, Sahara Y, Kitamura K (2015) Developmental increase in hyperpolarization-activated current regulates intrinsic firing properties in rat vestibular ganglion cells. Neuroscience 284:632–642.

Zhang H, Craciun LC, Mirshahi T, Rohács T, Lopes CMB, Jin T, Logothetis DE (2003) PIP2 Activates KCNQ Channels, and Its Hydrolysis Underlies Receptor-Mediated Inhibition of M Currents. Neuron 37:963–975.

